# Plant protoplast-based assay to screen for salicylic acid response-modulating bacteria

**DOI:** 10.1101/2022.11.02.514867

**Authors:** Moritz Miebach, Renji Jiang, Paula E. Jameson, Mitja N.P. Remus-Emsermann

## Abstract

Leaves host remarkably diverse microbes, collectively referred to as the leaf microbiota. While many beneficial functions have been attributed to the plant microbiota, the individual contributions of leaf-colonising bacteria range from pathogenic to mutualistic interactions. Omics approaches demonstrated that some leaf-colonising bacteria evoke substantial changes in gene expression and metabolic profiles in the plant host, including plant immunity. While omic approaches provide a system level view on cellular functions, they are costly and laborious, thereby severely limiting the throughput of the number of bacterial strains that can be tested *in planta*. To enable cost-effective high-throughput screens, we have developed a plant protoplast-based assay to measure real-time target gene expression changes following bacterial inoculation. Here, protoplasts were isolated from leaves of stable transgenic plants containing a pPR1:eYFP-nls construct. Changes in yellow fluorescence were captured for up to 96 treatments using a plate reader. This allowed the monitoring of changes in the salicylic acid-dependent plant immune response over time. Protoplast isolation per se evoked mild fluorescence responses, likely linked to endogenous salicylic acid production. This is advantageous in a bacterial assay, as bidirectional changes in PR1 expression can be measured. Plate reader-generated data were validated via fluorescence microscopy and RT-qPCR. Fluorescence microscopy further demonstrated heterogeneity in the response of individual protoplasts, which is potentially linked to differences in cell-type. In summary, the protoplast assay is an affordable and easily up-scalable way of measuring changes in target gene expression to bacterial colonisation.

## INTRODUCTION

Leaves are colonised by a vast diversity of bacteria, collectively referred to as the leaf microbiota. Large-scale isolations of bacteria from organ-specific microbiota enable bottom-up experiments (Bai et al. 2015; Vorholt et al. 2017). This allows the investigation of the individual contributions of plant-colonising bacteria to various aspects of plant life, including their influence on plant immunity. Some of these bacteria evoke gene expression and metabolic changes in the plant (Vogel et al. 2016; Ryffel et al. 2016). The magnitude of these changes differs depending on the leaf-colonising bacteria (Maier et al. 2021). While *in planta* transcriptomic and metabolomic studies provide highly resolved insights into the response of a plant to a specific leaf coloniser, these methods are too costly and time-intensive to investigate more than a handful of selected bacteria. Consequently, such studies can only represent a small fraction of the microbiome. Leaves, however, are colonised by many diverse bacteria, and their individual effects on the plant remain unknown. Thus, high throughput screening methods are badly needed.

Plants perceive microbes by highly conserved microbe-associated molecular patterns (MAMPs). MAMP perception triggers basal immunity, so-called pattern-triggered immunity (PTI) (Schlechter et al. 2019). An active immune system protects the plant from potential pathogen threat, but persistent immune activation infers a growth penalty (He et al. 2022). As MAMPs are present in microbes irrespective of their symbiotic relationship with the plant, the question arose as to whether and how plants differentiate between microbial colonisers with varying effects on the plant. Recently, it was shown that the beneficial rhizobacterium *Pseudomonas simiae* WCS417 elicited only half of the transcriptional responses than its cognate flg22 peptide. The suppressed genes were enriched in defence-related transcriptional responses. This suggests, either a suppression in MAMP recognition by the rhizobacterium, or an integration of multiple inputs, resulting in a weaker defence output in the plant (Stringlis et al. 2018). Such immune suppression appears to be highly prevalent among root-colonising bacteria. Out of 151 tested root-colonising bacteria 41% suppressed MAMP-triggered root growth inhibition (Ma et al. 2021). The suppression of immune responses likely limits costly immune responses in situations of no impending danger. Contrary to immune suppression, some non-pathogenic bacteria were shown to elicit mild immune responses in plants. The activation of plant immunity in these cases was potentially linked to the *in planta* protective ability of these non-pathogenic bacteria against specific foliar pathogens (Ritpitakphong et al. 2016; Vogel et al. 2016).

To investigate the effect of hundreds of individual leaf-colonising bacteria on plant immunity, a novel plant protoplast-based assay was developed, and its utility is described here. Plant protoplasts have been isolated for more than 60 years (Cocking 1960) and are commonly used for transient gene expression in conjunction with microscopy (Yanagisawa, Yoo, and Sheen 2003; Cho, Yoo, and Sheen 2006; Yoo, Cho, and Sheen 2007; Wu et al. 2009) or cell sorting for cell-type specific omics studies (Birnbaum et al. 2003; Ryu et al. 2019). In this study, a stable transgenic plant line containing a pPR1:eYFP-nls construct (Betsuyaku et al. 2018) was employed to determine changes in Pathogenesis-Related Gene 1 (PR1) expression by measuring enhanced Yellow Fluorescent Protein (eYFP) fluorescence. PR1 is a robust marker for salicylic acid (SA) pathway activity (Vlot, Dempsey, and Klessig 2009; Pieterse et al. 2012), which itself is central to plant immunity (Gaffney et al. 1993; Cao et al. 1994; Delaney, Friedrich, and Ryals 1995; Nawrath and Métraux 1999), and induced by various plant-associated bacteria (Vogel et al. 2016; Shi et al. 2022).

## MATERIAL AND METHODS

### Plant material and growth

Experiments were conducted on *Arabidopsis thaliana* wild-type (Col-0) and pPR1:eYFP-nls transgenic plants. The pPR1:eYFP-nls transgenic line was kindly provided by A/Prof. Shigeyuki Betsuyaku (University of Tsukuba, Japan).

Seeds were surface-sterilised according to Lundberg et al. (2012) and sown on cut pipette tips (200 µL) filled with ½ MS 1% phytoagar, pH 5.9. Seven days after sowing, the pipette tips holding the seedlings were transferred to autoclaved plant tissue culture boxes (Magenta vessel GA-7, Magenta LLC, Lockport, IL, USA), filled with 100 ml of ½ MS 1% phytoagar at pH 5.9. Four seedlings were transferred per box. Lids of the plant tissue culture boxes contained four holes for gas exchange (9 mm diameter), which were covered by two pieces of micropore tape (3M, Saint Paul, USA) to ensure axenic conditions. Plant growth and seedling germination took place in a CMP6010 growth cabinet (Conviron, Winnipeg, Canada) with 11 h of light (150–200 µmol m^−2^ s^−1^) and 85% relative humidity.

### Mesophyll protoplast isolation

Leaf protoplasts were isolated as described in Yoo et al., and Wu et al., with slight modifications (Yoo, Cho, and Sheen 2007; Wu et al. 2009). The enzyme solution 1% (w/v) cellulase R10 (Duchefa, Haarlem, Netherlands), 0.25% (w/v) macerozyme R10 (Duchefa, Haarlem, Netherlands), 100 mM 2-ethanesulfonic acid (MES, pH 5.7; Duchefa, Haarlem, Netherlands), 20 mM KCl, 10 mM CaCl_2_, 0.1% (w/v) BSA was freshly prepared prior to protoplast isolation. BSA and CaCl_2_ were added after the solution was kept at 55°C for 10 min and cooled down to room temperature. The enzyme solution was filter-sterilised into a petri dish with a 0.2 µm syringe filter. Strips of autoclave tape (3M, Saint Paul, USA) and masking tape (Dixon, New Zealand) were UV-sterilised for 15 min in a biological safety hood. Mature leaves from healthy, axenic six- to eight-weeks-old plants were collected with sterile scissors and tweezers and fixed onto the autoclave tape with the adaxial side. The masking tape was then firmly pressed onto the abaxial side of the leaves using the bottom of a microcentrifuge tube (MCT-150-C, Axygen, Corning, USA). The masking tape was then carefully peeled away. The cuticle of the abaxial leaf had adhered to the masking tape. The leaves were then submerged in 10 ml enzyme solution. The samples were incubated at room temperature and constant orbital shaking at 40 rounds per minute for 60-90 min. Next, the solution was carefully transferred via pipetting into a 50 ml falcon tube, followed by centrifugation at 100× *g* for 3 min. The supernatant was discarded, and the protoplast pellet washed twice with 20 ml of pre-chilled W5 buffer (154 mM NaCl, 125 mM CaCl_2_, 5 mM KCl, 5 mM glucose, 2 mM MES (pH 5.7)) followed by centrifugation at 100× *g* for 3 min. The protoplast solution was kept at 4°C, while its concentration was determined with a hemocytometer (Neubauer, Germany). Protoplast density was adjusted to 10^5^ cells ml^-1^ and immediately used for downstream experiments.

### Protoplast assay

Protoplasts isolated from WT and pPR1:eYFP-nls plants were distributed in a 96-well plate (Corning, Corning, USA). To that end, 100 µl of protoplast solution from either plant line was mixed with 100 µl of salicylic acid (SA) solution of differing concentrations (0.005 - 0.5 mM) or 100 µl of bacterial suspension (Table 1). SA was dissolved in the W5 buffer. Bacteria were cultivated at 30°C on R2A (HIMEDIA LABORATORIES, Mumbai, India) media plates. Bacterial suspensions were prepared from bacterial colonies suspended in the W5 buffer and washed twice via centrifugation at 4000× *g* for 5 min followed by discarding the supernatant and suspension in W5. The bacterial suspensions were then adjusted to ∼ 2 × 10^7^ cfu ml^-1^.

**Table 1:** List of bacterial strains used in this study.

| Bacterial strain | Reference | cfu ml<sup>-1</sup> at<br>OD<sub>600</sub> = 1 |
| --- | --- | --- |
| *Aeromicrobium* sp. Leaf245 | (Bai et al. 2015) | $2 \times 10^9$ |
| *Methylobacterium radiotolerans* 0-1 | (Kwak et al. 2014) | $1.1 \times 10^8$ |
| *Methylobacterium* sp. Leaf92 | (Bai et al. 2015) | $6.1 \times 10^7$ |
| *Pseudomonas citronellolis* P3B5 | (Remus-Emsermann et al. 2016) | $1 \times 10^9$ |
| *Pseudomonas koreensis* P19E3 | (Schmid et al. 2018) | $1 \times 10^9$ |
| *Rhodococcus* sp. Leaf225 | (Bai et al. 2015) | $9.1 \times 10^7$ |
| *Sphingomonas phyllosphaerae* FA2 | (Rivas et al. 2004) | $6.7 \times 10^8$ |
| *Sphingomonas* sp. Leaf17 | (Bai et al. 2015) | $6.7 \times 10^8$ |
| *Sphingomonas* sp. Leaf357 | (Bai et al. 2015) | $2.9 \times 10^8$ |
| *Williamsia* sp. Leaf354 | (Bai et al. 2015) | $1.5 \times 10^8$ |

The fluorescence signal of the protoplast response to SA was measured in a plate reader (BMG Labtech, Ortenberg, Germany). Every 5 min, yellow fluorescence (Ex.485/15, Em.520/12) was measured at a gain of 2000 with 160 flashes per well at a diameter of 3 mm per well. Individual experiments were run for up to 24 h at a temperature of 25°C. The data generated by the plate reader were further processed and analysed in the R programming environment (R Core Team 2021). Baseline fluorescence was estimated per well by averaging the fluorescence intensity measurements within the first hour after treatment and subtracting this from the remaining data. Next, a cubic smoothing spline was added to the data using the ‘smooth.spline’ function with the number of knots set to four (Chambers and Hastie 1992). To calculate the area under the curve (AUC), first each curve was raised by the absolute value of the lowest value of all curves of a given plant type per run, to eliminate negative AUCs in each curve. The AUC was calculated using the composite trapezoid rule via the ‘auc’ function from the ‘MESS’ package (Ekstrøm 2020). Next the AUCs were normalised against the mean AUC of the mock (buffer only) controls per plant type.

### Microscopy and image processing

The fluorescence signal of the protoplast response to SA was measured 17 h after treatment with SA. Microscopy was performed using an Olympus IX70 fluorescence microscope (Olympus, Shinjuku City, Japan) at 200× magnification (Objective Olympus LCPlanFl 20× NA 0.4) equipped with Leica filter set U-MWU (excitation bandpass 330-385 nm, dichroic mirror 400 nm, emission long pass filter 420 nm) for the detection of eYFP. Images were acquired with a AxioCam HRc (Zeiss, Jena, Germany) in conjunction with the software AxioVision 4 (version 4.8.2.0) at 10 ms for quantitative and 200 ms exposure time for qualitative image analysis and a digital gain of 4. Brightfield images were taken at 6 ms exposure time. Image processing was performed in FIJI/ImageJ (Version 2.1.0/1.53c) (Schindelin et al. 2012). Per image the outlines of 30 randomly selected, but intact looking, protoplasts were manually traced based on the brightfield channel. After a ‘rolling ball’ background subtraction with a radius of 50 pixels, the mean and maximal YFP fluorescence intensity of every traced protoplast was determined. The percentage of eYFP expressing protoplasts in the transgenic line was determined as the number of protoplasts with a maximal fluorescence above the maximal fluorescence (pixel intensity > 13) in all measured wild-type protoplasts.

### Gene expression analysis

Protoplasts were sampled 17 h after treatment with SA. Two biological replicates were sampled per SA concentration. RNA extraction, cDNA synthesis and RT-qPCR was performed as previously described in Miebach et al. (2020). Primers that were used in this study are listed in Table 2.

**Table 2:**
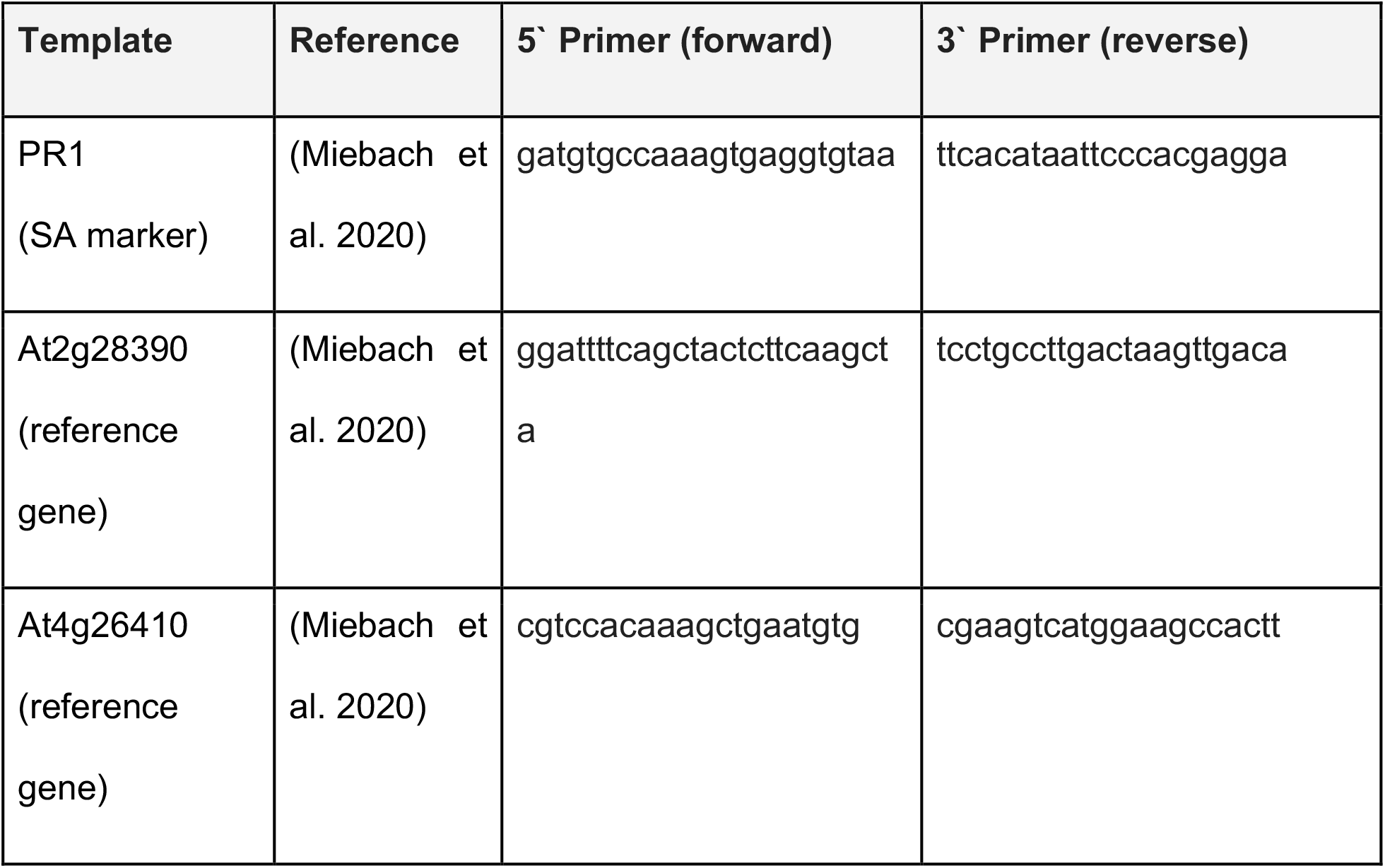
List of primers used in this study.

## RESULTS

Protoplasts were isolated under sterile conditions from leaves of seven-weeks-old arabidopsis wild-type (Col-0) and transgenic (pPR1:eYFP-nls) plants, that were grown axenically in the ‘Litterbox’ system, as described in Miebach et al. (2020). The protoplasts were isolated using the cuticle tape lift method (Wu et al. 2009). Fresh protoplasts were distributed in a 96-well plate to reach approximately 10^4^ protoplasts in 200 µL per well. To determine the threshold of the assay the protoplasts were treated with different SA concentrations, to elicit the expression of eYFP-nls driven by the PR1 promoter (Fig. 1).

**Fig. 1.**
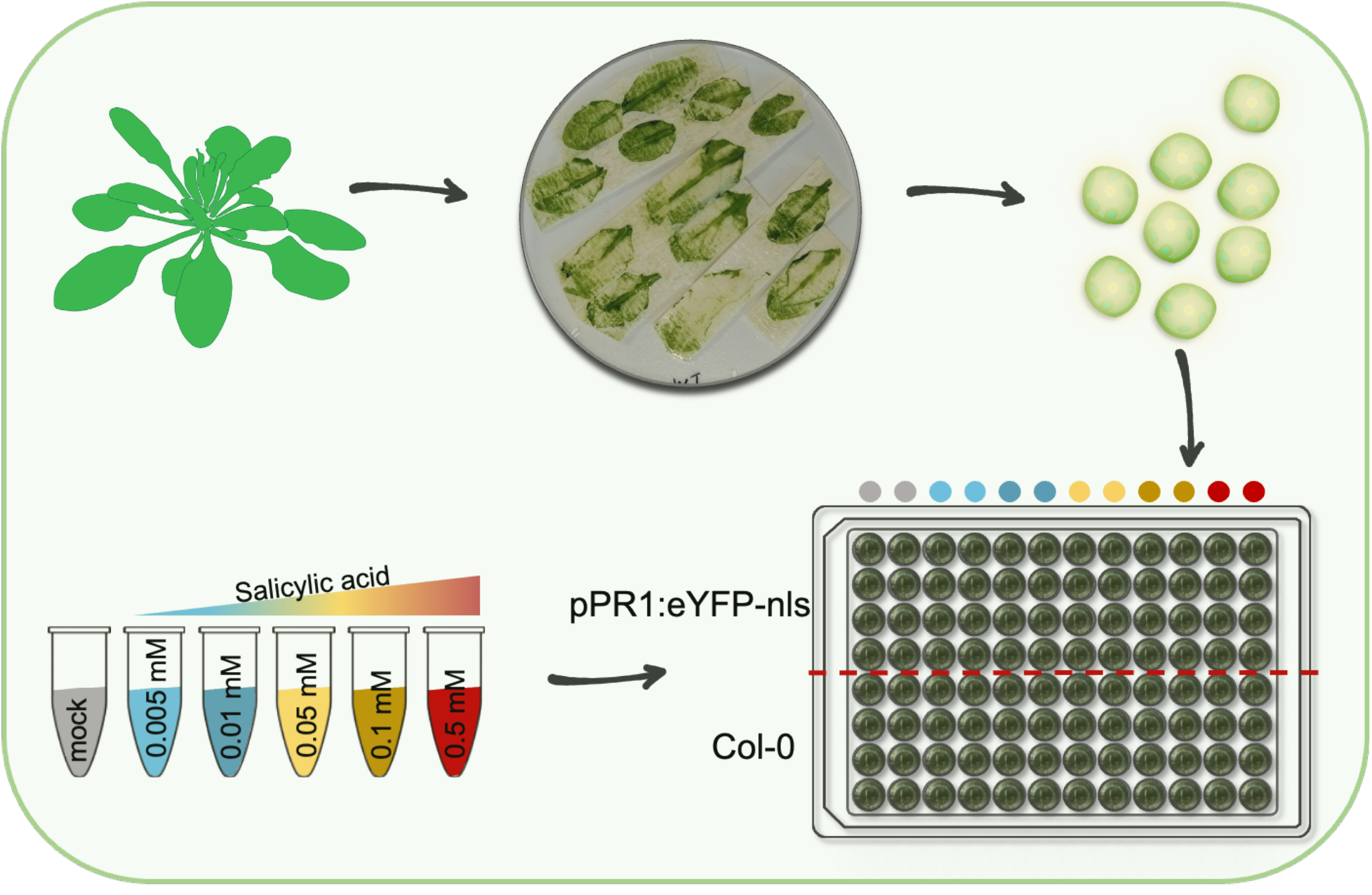
Experimental design. Protoplasts were isolated from leaves of axenic seven-weeks-old arabidopsis wild-type (Col-0) and transgenic (pPR1:eYFP-nls) plants using the cuticle tape lift method (Wu et al. 2009). The top centre circle shows the remaining plant leaf material after enzymatic digestion for 1-1.5 h. Approximately 10^4^ protoplasts per well were distributed in 96 well plates and treated with different SA concentrations, as indicated by the coloured dots above the 96-well plate.

Yellow fluorescence was measured over time in a plate reader for up to 24 h. Response curves were generated from background-subtracted and spline-fitted fluorescence intensity measures (Fig. 2). Under most conditions, fluorescence dropped to a minimum at around 5 h post treatment (Fig. 3A and B). Interestingly, this drop was observed in both wild-type and transgenic protoplasts, indicating that it was unrelated to eYFP expression.

**Fig. 2.**
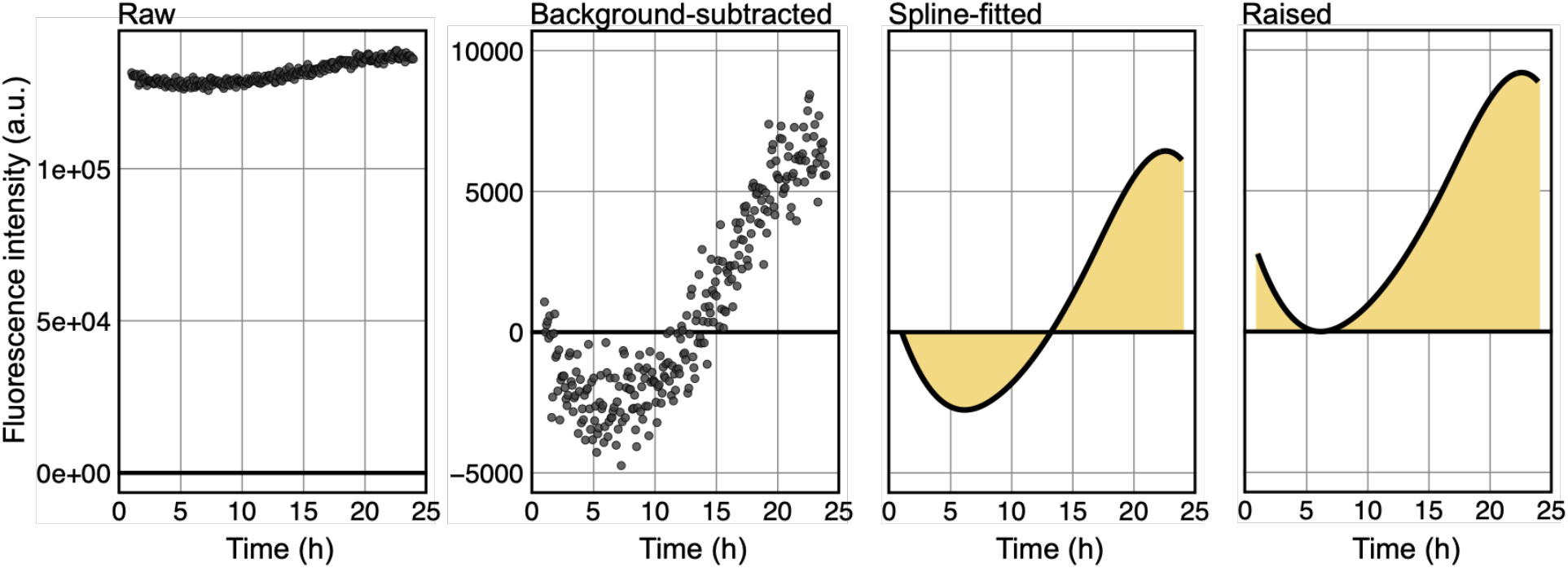
Fluorescence response curve data processing pipeline. Background fluorescence was estimated from fluorescence reads from within the first hour post treatment. The estimated background fluorescence was then subtracted from raw fluorescence reads and a spline curve was fitted to the reads. The spline curves were raised by a set value per plant type (Col-0; pPR1:eYFP-nls) to then estimate the correct area under the curve.

**Fig. 3.**
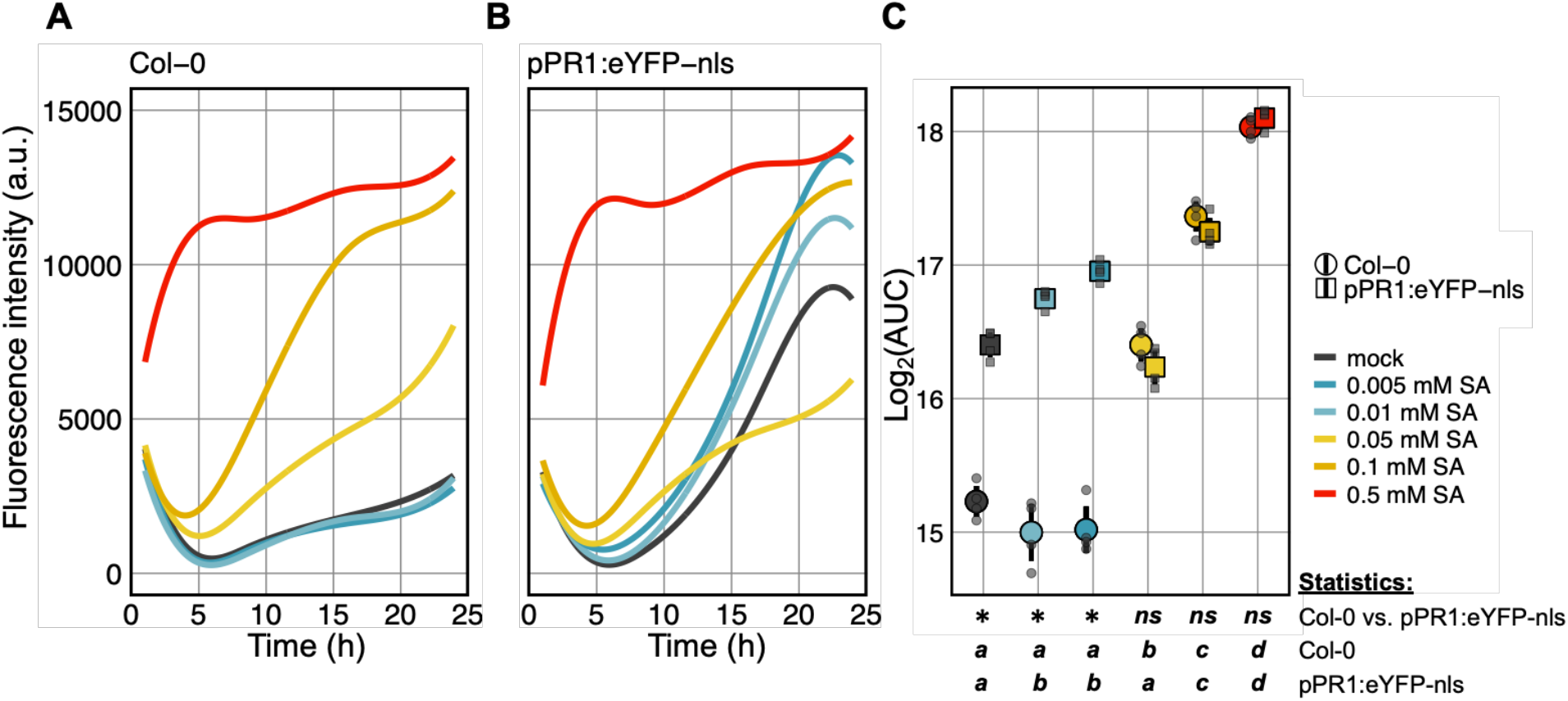
Protoplast response to different SA concentrations as determined by a fluorescent plate reader. A,B. Yellow fluorescence over 24 h in response to different concentrations of SA in wild-type (A) and transgenic (B) protoplasts. C. Data depicted in A,B expressed as area under the curve (AUC). Coloured shapes depict the mean, bars depict the standard error, colour depicts SA concentration, grey shapes depict the AUC from individual replicates, * depicts statistical significance (*p* < 0.05), *ns* depicts the lack of statistical significance in fluorescence expression between wild-type and transgenic protoplasts, Mann-Whitney Test. Letters depict statistical differences (*p* < 0.01) in protoplast response to different SA concentrations for wild-type and transgenic protoplasts, one-way ANOVA & Tukey’s HSD test. Shown are four technical replicates per plant type and SA concentration.

### Protoplasts deteriorated at high concentrations of SA

At SA concentrations ≥ 0.05 mM, a strong fluorescence response was observed in wild-type protoplasts, suggesting that a physiological change in these protoplasts led to autofluorescence in parts of the eYFP spectrum (Fig. 3A). In addition, these response curves closely mimicked the response curves observed in transgenic protoplasts at these concentrations of SA, thereby suggesting that no quantifiable amounts of eYFP were expressed in the transgenic protoplasts (Fig. 3B and C). Further, the response curves increased in intensity with increasing SA concentrations. This indicates that the physiological change in the protoplasts was SA concentration-dependent at concentrations ranging from 0.05 mM to 0.5 mM. Brightfield microscopy further confirmed that protoplast health deteriorated at SA concentrations ≥ 0.05 mM (Fig. 4A-F). Protoplasts decreased in size, darkened and their plasma membrane retracted with increasing SA concentration, indicating the loss of cell membrane integrity and vacuole rupture, both hallmarks of programmed cell death (Fig. 4D-F) (Young et al. 2010).

**Fig. 4.**
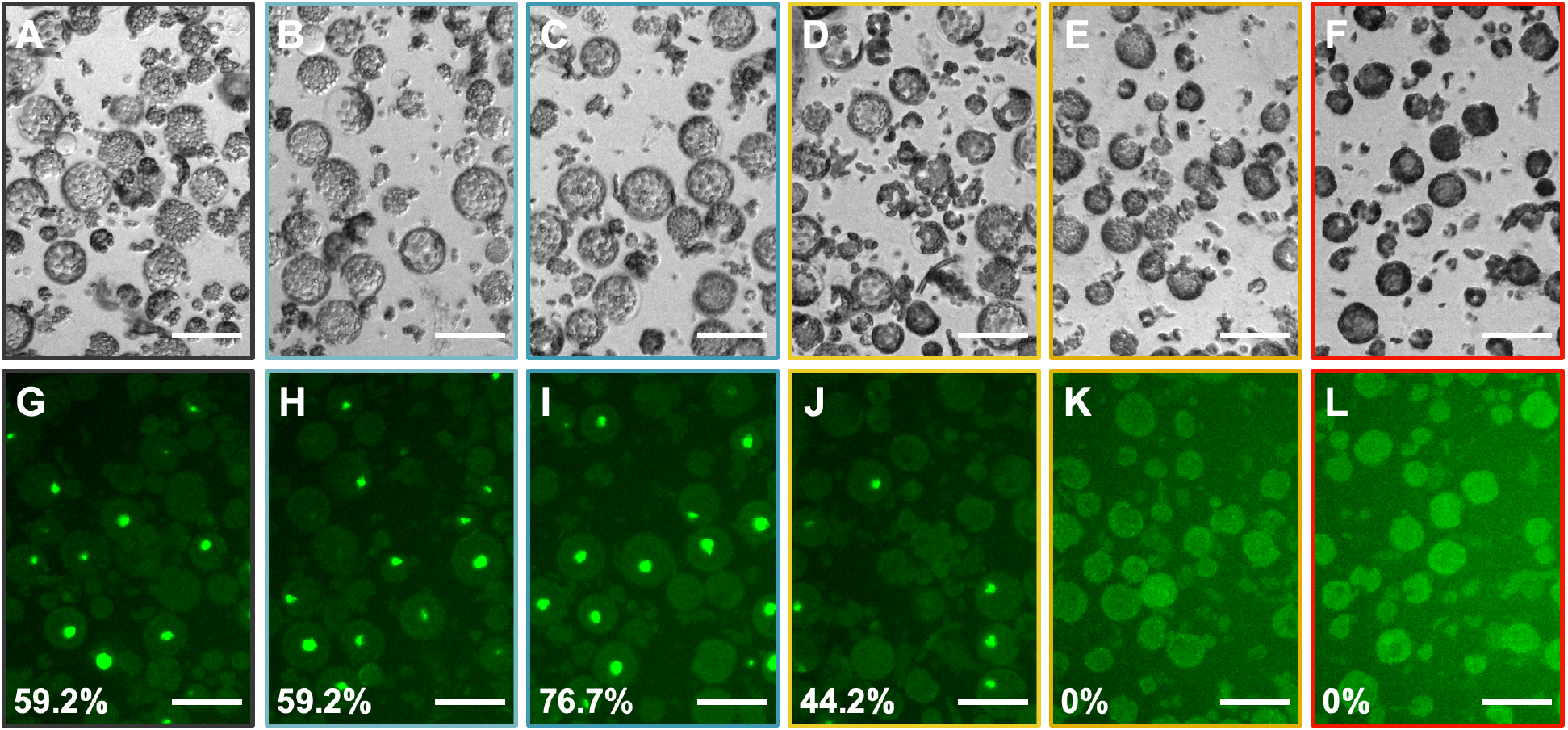
Individual protoplast response to different SA concentrations. A-L. Brightfield (A-F) and fluorescence (G-L) images of protoplasts treated with 0 (A,G), 0.005 (B,H), 0.01 (C,I), 0.05 (D,J), 0.1 (E,K), or 0.5 (F,L) mM SA. Bottom left of each fluorescence image (G-L) depicts the percentage of eYFP-nls expressing protoplasts. n = 120 per SA concentration. Bar scale 100 µm.

### The expression of eYFP was SA concentration-dependent

At SA concentrations ≤ 0.01 mM only a weak response curve was observed in wild-type protoplasts. Importantly, the curves were indistinguishable from those of mock-treated protoplasts (Fig. 3A). In contrast, strong response curves were observed at these concentrations of SA in transgenic protoplasts, suggesting the expression of eYFP. Their maximum fluorescence peaked at around 22 h post treatment and with increasing SA concentration the maximum fluorescence increased (Fig. 3B) as well as the area under the curve (AUC) (Fig. 3C). Interestingly, the mock control itself exhibited a strong response curve in transgenic protoplasts that was clearly distinguishable from the fluorescence response in mock-treated wild-type protoplasts (Fig. 3A and B). The curve followed the shape of those at SA concentrations ≤ 0.01 mM, but slightly weaker in the overall response (Fig. 3B and C). This suggests that the state of being a protoplast itself elicits the expression of PR1 and thereby eYFP in the transgenic line.

### Heterogeneity in the response of individual protoplasts

Fluorescence microscopy revealed heterogeneity in the presence of nuclear-localised eYFP in individual protoplasts 17 h post treatment (Fig. 4G-J; Fig. 5B). The percentage of eYFP-nls expressing protoplasts depended on the SA concentration. An increase in SA-responsive protoplasts from 59.2% in mock-treated and 0.005 mM SA-treated to 76.7% in 0.01 mM SA-treated protoplasts was observed (Fig. 4G-I; *p* < 0.001, Fisher exact test). Interestingly, even though no significant difference in the AUC of wild-type and transgenic protoplasts was observed at 0.05 mM SA, still 44.2% of the measured protoplasts were expressing eYFP-nls (Fig. 3C; Fig. 4J). This indicates that the slight expression of eYFP-nls at 0.05 mM SA was masked by the induced protoplast autofluorescence. At SA concentrations ≥ 0.1 mM no nuclear localised fluorescence was detected. Instead, the protoplasts exhibited a general autofluorescence (Fig. 4K and L).

**Fig. 5.**
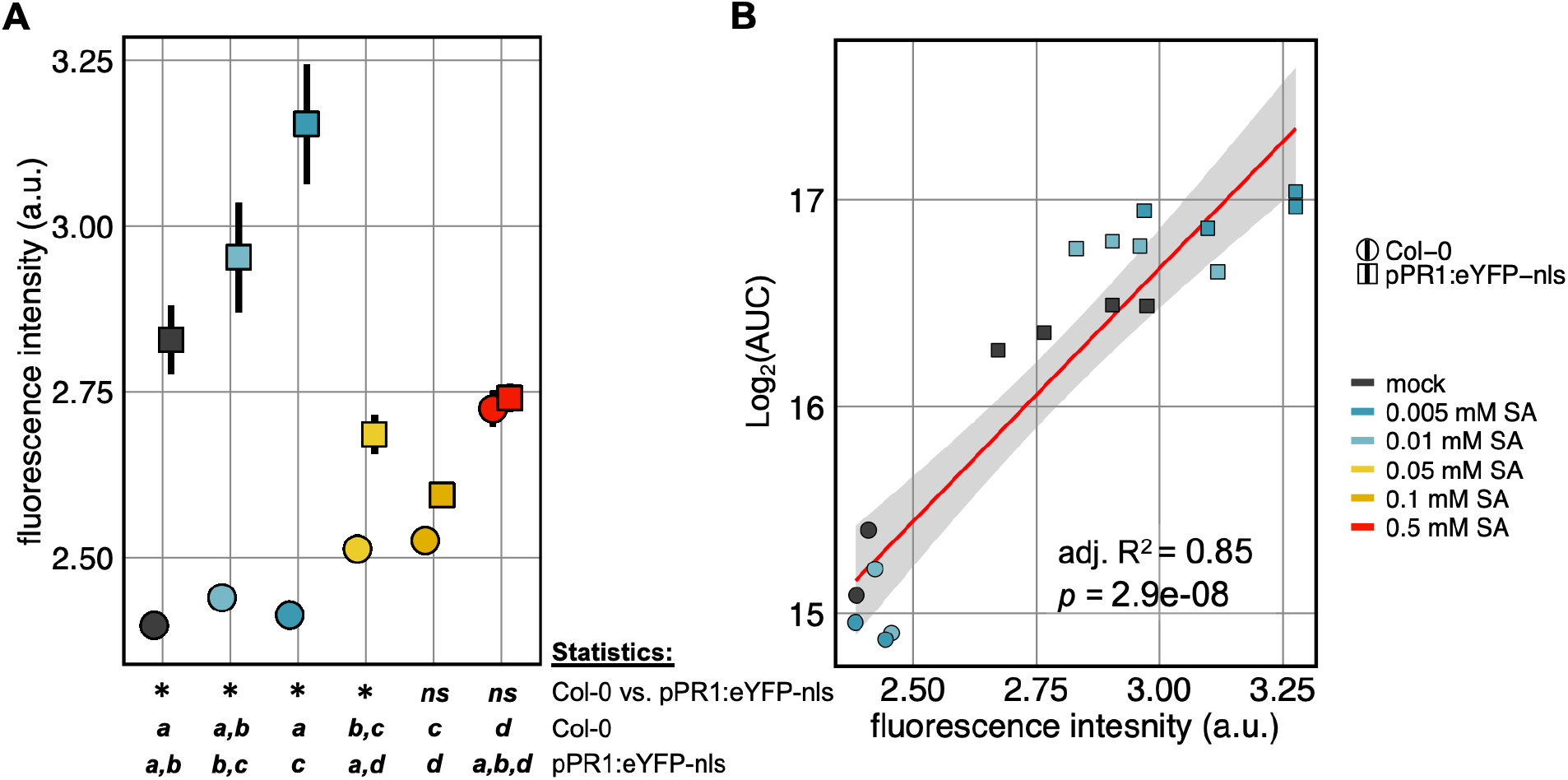
Fluorescence intensity of individual protoplasts in response to different SA concentrations as determined by fluorescence microscopy. A. Protoplast eYFP fluorescence 17 h post treatment. Shapes depict the mean, bars depict the standard error, colour depicts SA concentration, * depicts statistical significance (*p* < 0.05), *ns* depicts the lack of statistical significance in fluorescence expression between wild-type and transgenic protoplasts, Mann-Whitney Test. Letters depict statistical differences (*p* < 0.05) in protoplast response to different SA concentrations for wild-type and transgenic protoplasts, one-way ANOVA & Tukey’s HSD test. n = 120 per SA concentration in pPR1:eYFP-nls, n = 60 per SA concentration in Col-0. B. Correlation between fluorescence intensity measured from microscopy images and area under the curve (AUC) from the plate reader experiment (see Fig. 3). Shapes depict the mean, colour depicts SA concentration, red line depicts a fitted linear model, the grey bar depicts limits of 95% confidence interval.

### Validation of plate reader results by independent methods

Fluorescent microscopy further validated that the dose-dependent signal at SA concentrations ≤ 0.01 mM observed in the plate reader assay was indeed largely related to the expression of eYFP-nls (Fig. 3C; Fig. 5A). At SA concentrations ≤ 0.01 mM 85% (adj. R^2^ = 0.85, *p* = 2.9 × 10^−8^) of the response of the protoplasts in the plate reader assay was explained by the individual fluorescence response of protoplasts measured at 17 h post treatment (Fig. 5B).

The gene expression levels of PR1, as determined by RT-qPCR, further validated that the measured eYFP fluorescence was indeed related to PR1 expression (Fig. 6). At SA concentrations ≤ 0.01 mM an up to 4-fold increase in PR1 mRNA levels in protoplasts was observed. Further, it should be noted that at SA concentrations ≥ 0.1 mM no RNA of sufficient quality could be extracted, again suggesting that the protoplasts were deteriorating at these concentrations.

**Fig. 6.**
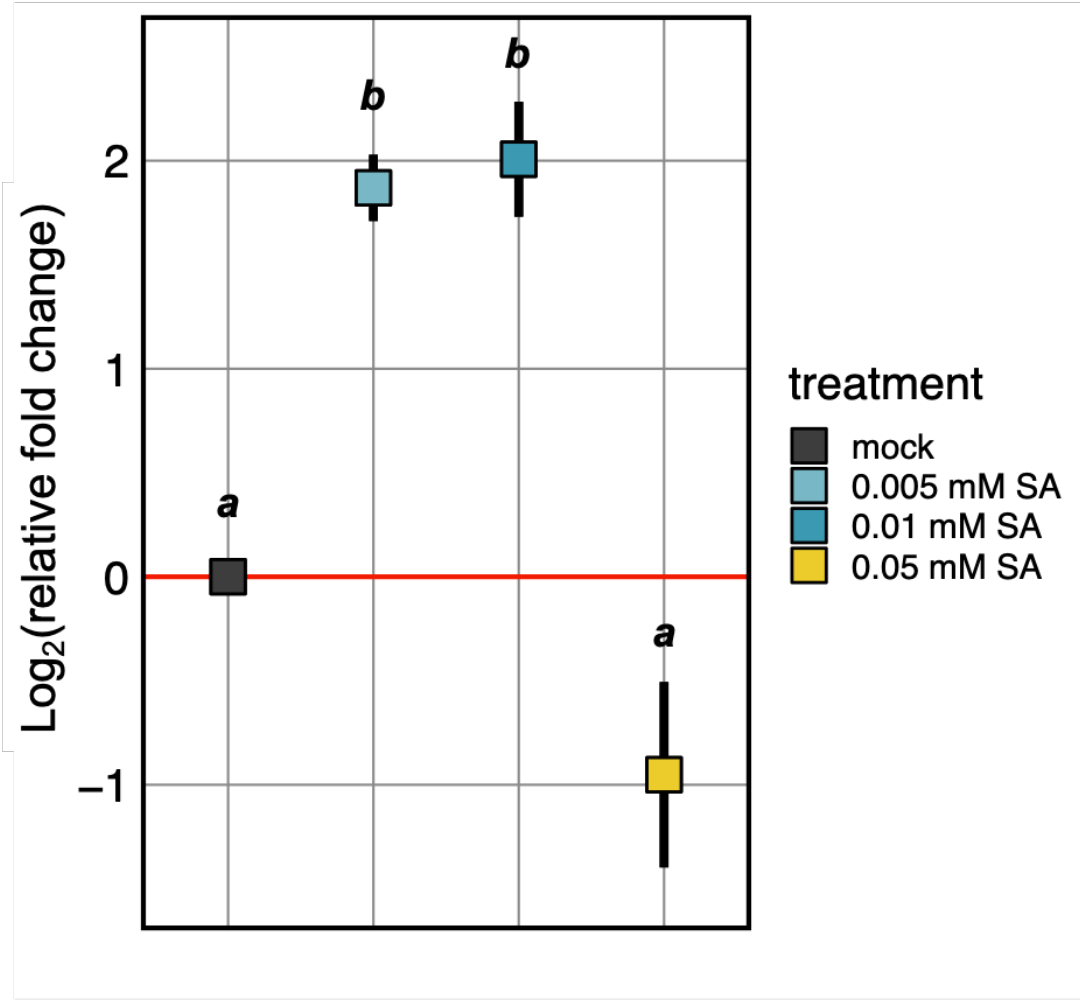
PR1 expression in protoplasts in response to different SA concentrations. Log_2_ fold-change in gene expression relative to mock-treated control 17 h post treatment. Squares depict the mean, bars depict the standard error, colour depicts SA concentration. Letters depict statistical differences (*p* < 0.05) in protoplast response to different SA concentrations, one-way ANOVA & Tukey’s HSD test. n = 2 biological replicates with technical triplicates per biological replicate.

### Protoplasts isolated from young plants tolerate higher SA concentrations

To test whether plant age influenced the responsiveness of the protoplasts to exogenous SA, protoplasts were simultaneously isolated from three different aged plant batches and treated with different concentrations of SA. Compared with the previous experiments, the range of SA concentrations was slightly narrowed, in order to obtain a better overview of the threshold concentration of SA at which autofluorescence becomes detectable, a hallmark for the onset of protoplast deterioration. Protoplasts isolated from eight-weeks-old wild-type plants exhibited significant increases in autofluorescence already at concentrations as low as 0.01125 mM SA (Fig. 7). Therefore, they seemed to tolerate less SA than those isolated from younger plants, which showed no significant increase in autofluorescence at SA concentrations ≤ 0.0225 mM (Fig. 7). The response of transgenic protoplasts appeared to be more consistent throughout the different age groups, but with a slight decrease in overall fluorescence in protoplasts of eight-weeks-old plants.

**Fig. 7.**
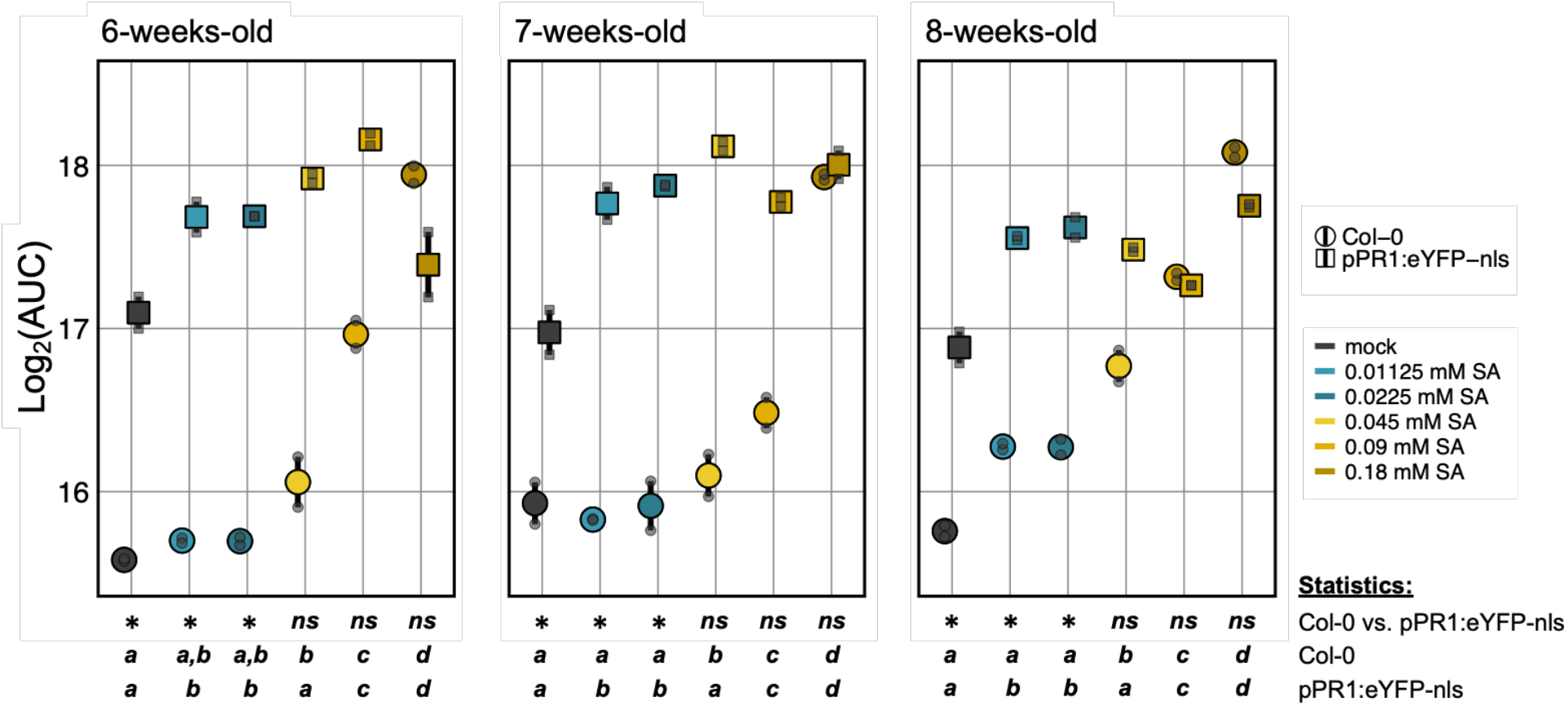
Protoplast response to different SA concentrations in relation to plant age. AUC of protoplasts from different aged plants (six-, seven- and eight-weeks-old plants) and different plant types. Coloured shapes depict the mean, bars depict the standard error, colour depicts SA concentration, grey shapes depict the AUC from individual replicates. Letters depict statistical differences (*p* < 0.05) in protoplast response to different SA concentrations for wild-type and transgenic protoplasts, one-way ANOVA & Tukey’s HSD test. Shown are two technical replicates per plant type, plant age and SA concentration.

### Some non-pathogenic bacteria elicit an eYFP response in leaf protoplasts

To demonstrate the application of the protoplast assay, protoplasts were inoculated with various non-pathogenic leaf-colonising bacterial strains. Four out of ten strains exhibited a peak fluorescence at least 1.4× higher than the mock-treated control (Fig. 8B). *Sphingomonas* sp. Leaf17 elicited the strongest response in the protoplasts, followed by *Pseudomonas citronellolis* P3B5, *Pseudomonas koreensis* P19E3 and *Methylobacterium* sp. Leaf92. The protoplast response to members of the family Nocardioidaceae was either relatively weak (*Rhodococcus* sp. Leaf225 and *Williamsia* sp. Leaf354) or non-existent (*Aeromicrobium* sp. Leaf245). Interestingly, in the case of Sphingomonodaceae and Methanobacteriaceae the strains that were isolated from arabidopsis (*Sphingomonas* sp. Leaf17, *Sphingomonas* sp. Leaf357 and *Methylobacterium* sp. Leaf92) did elicit eYFP expression in protoplasts, whereas strains isolated from leaves of different hosts (*Sphingomonas phyllosphaerea* FA2 and *Methylobacterium radiotolerans* 0-1) did not (Ito and Iizuka 1971; Green and Bousfield 1983; Bai et al. 2015).

**Fig. 8.**
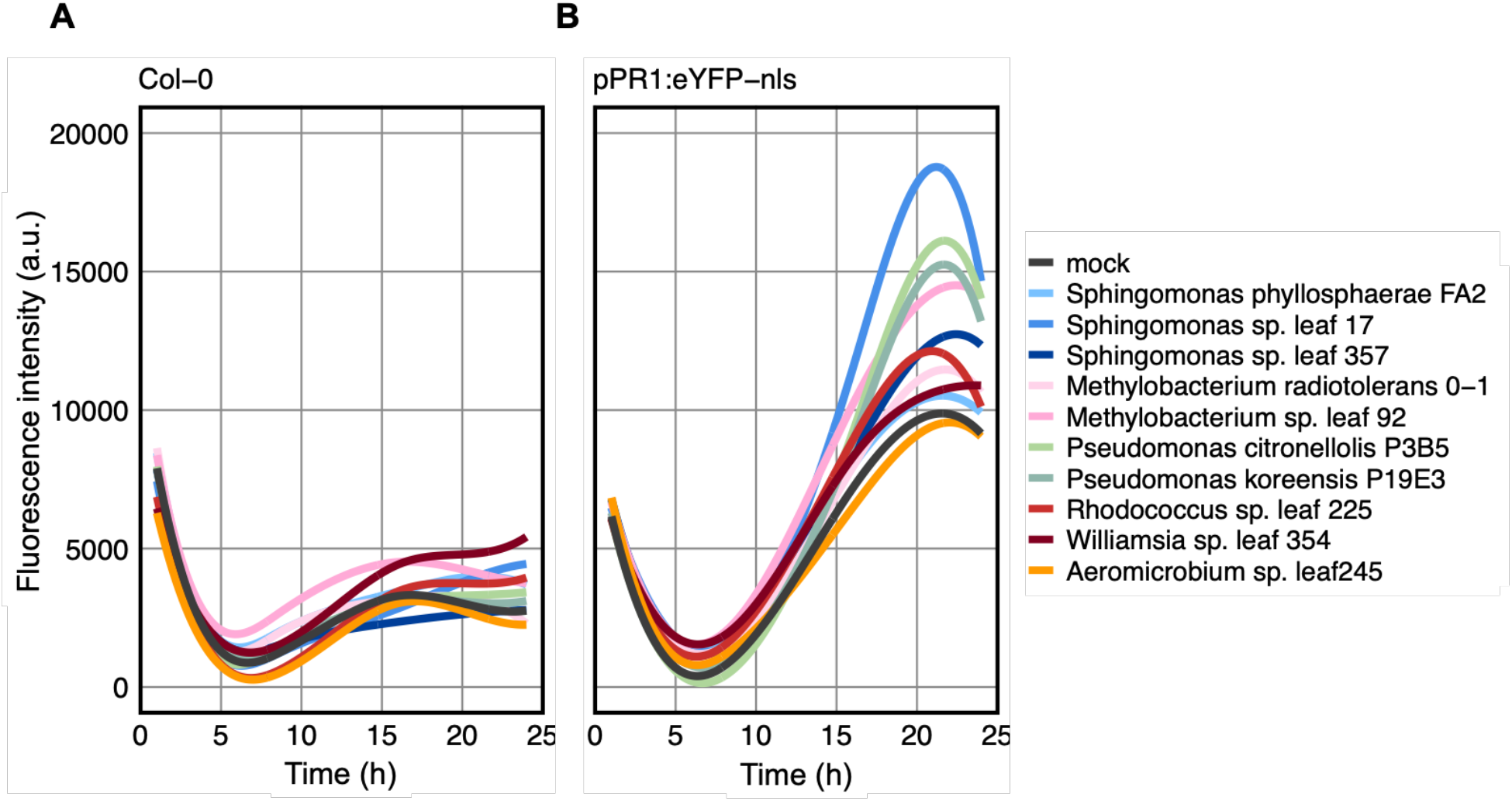
Protoplast response to different bacteria as determined by a fluorescent plate reader. A,B. Yellow fluorescence over 24 h in response to different bacteria in wild-type (A) and transgenic (B) protoplasts. Shown are the mean of two technical replicates per plant type and bacterial inoculant. Bacterial inoculants are sorted by phylogeny (Oso et al. 2019).

The expression of yellow fluorescence in wild-type protoplasts varied between treatments (Fig. 8A). However, the observed expression profiles were not comparable to those observed in wild-type protoplasts treated with SA at concentrations ≤ 0.05 mM (Fig. 3A, Fig. 8A). Further, these mild changes cannot explain the strong changes that were observed in the transgenic line.

## DISCUSSION

### The protoplast assay monitors bidirectional changes in PR1 expression

We have shown that the protoplast assay can be employed to monitor changes in PR1 expression. At SA concentrations ≤ 0.01 mM an increase in protoplast fluorescence was observed, which was solely attributed to eYFP expression, as wild-type protoplasts showed no differences in fluorescence (Fig. 3). Further, up to a 4-fold change in PR1 expression was observed at concentrations ≤ 0.01 mM of exogenous SA in comparison to mock-treated control (Fig. 6). It should be noted that the mock control also expressed eYFP, likely driven by endogenous SA levels induced by the protoplastation (Fig. 3). Therefore, the assay can monitor up to ∼ 12-fold changes in PR1 expression, considering (1) that a 4-fold increase in PR1 gene expression represents a ∼ 1.4-fold increase in AUC, between mock-treated and 0.01 mM SA-treated protoplasts (Fig. 3C, Fig. 6), and (2) that a 4-fold change in AUC was observed between wild-type and transgenic protoplasts treated with 0.01 mM SA (Fig. 3C). Changes within this range of PR1 expression were previously reported upon leaf inoculation with two different bacteria (Vogel et al. 2016). However, how transferable the data are between an *in vitro* protoplast assay and *in planta* experiments remains to be seen.

As previously mentioned, protoplastation itself led to the activation of the PR1 promoter, leading to a 2-fold change in the AUC between wild-type and transgenic mock-treated protoplasts (Fig. 3C), roughly corresponding to a ∼ 6-fold increase in PR1 expression. In the context of a bacterial assay this is beneficial as it allows the monitoring of bidirectional changes in PR1 expression. The suppression of plant immunity for example appears to be a common trait among many root-colonising bacteria, as was recently demonstrated indirectly by monitoring the suppression of plant immune related root-growth inhibition (Ma et al. 2021).

The average fluorescence response of individual protoplasts was strongly correlated with the plate reader results at SA concentrations ≤ 0.01 mM (Fig. 3C, Fig. 5A and B). Consequently, the protoplast assay can also be performed by fluorescence microscopy, which has the advantage of better spatial resolution. Indeed, it would be possible to perform these experiments using time-lapse fluorescence microscopy if a suitable automated microscope was at hand, thus simultaneously providing the temporal resolution of the plate reader and the spatial resolution of the microscope.

Theoretically, the protoplast assay can be extended to any target gene of choice. With regard to plant immunity there are other interesting targets to be tested in future studies, including VSP2, a common target to measure jasmonic-acid related immune outputs (Chini et al. 2007; Mousavi et al. 2013), and PDF1.2, a distinct marker for a combined input of jasmonic acid and ethylene signalling (Lorenzo et al. 2003; Pré et al. 2008; Zhu et al. 2011). Excellent plant lines for the investigation of these targets were recently developed (Ghareeb et al. 2020). These plant lines express a reference fluorophore next to a fluorescent marker that is driven by the target gene promoter. This is advantageous in the case of fluorescence microscopy, as nuclei can be automatically traced (Ghareeb et al. 2020).

### Protoplasts deteriorate at high concentrations of SA

At SA concentrations ≥ 0.05 mM a concentration-dependent decay in protoplast health was observed. Protoplasts exhibited a shrivelled morphology indicating the loss of cell membrane integrity and vacuole rupture, both hallmarks of programmed cell death (Young et al. 2010) (Fig. 4D-F). Further, a strong fluorescence response was observed in wild-type protoplasts in the plate reader assay (Fig. 3A and C), suggesting the expression of autofluorescent compounds in response to physiological changes. High SA levels are known to trigger the hypersensitivity response, a programmed cell death response that usually occurs at pathogen entry sites (Mur et al. 2008). In agreement with the findings in the current study, it was previously reported that high concentrations of SA (1 mM) lead to 90% cell death in tobacco cell suspensions within 24 h, whereas no cell death was observed at SA concentrations ≤ 0.1 mM (Norman et al. 2004). The suspended tobacco cells seem to tolerate higher concentrations of exogenously applied SA, than freshly isolated protoplasts from arabidopsis leaves. However, this might not relate to endogenous levels, as mock-treated tobacco cells did not contain any detectable levels of SA (Norman et al. 2004). In the current study, mock-treated protoplasts expressed eYFP-nls, suggesting the presence of SA. However, how comparable data are from tobacco suspension cells and freshly isolated arabidopsis protoplasts is as yet unknown. In addition, cells from different plant species might have different sensitivities to SA.

Fluorescence microscopy revealed that at 0.05 mM SA, the onset concentration of protoplast deterioration, 44.2% of the protoplasts expressed eYFP (Fig. 4J). At SA concentrations ≥ 0.1 mM no eYFP expression was detected (Fig. 4K and L). Whether eYFP was expressed at SA concentrations ≥ 0.1 mM before the protoplasts were investigated by fluorescence microscopy, 17 h post treatment, remains uncertain. During programmed cell death the cytosol acidifies, which then leads to a reversible attenuation of the YFP signal (Young et al. 2010).

### Heterogeneity in the protoplast response

Heterogeneity in nuclear localised eYFP expression of individual protoplasts was observed via fluorescence microscopy (Fig. 4). This heterogeneity within the protoplast population is likely driven by differences in the competence of individual protoplasts to respond to SA. Most of the protoplasts used in this assay likely originated from the mesophyll. Kim et al. (2021) reported 75% of their protoplasts used for single-cell RNA-seq were of mesophilic origin. However, even though the protoplast preparation protocol used by these workers was very similar to the one used in the current study, they cut the leaves around the major vein to enrich cells of vascular origin, thereby lowering the number of mesophyll cells (Kim et al. 2021). Further, they reported on 11 different cell clusters within the mesophyll cell population. An analysis of the gene expression of these cell clusters would be an interesting subject for future research to infer their epigenetic states and thus their competency to respond to SA.

Interestingly, at different SA concentrations the number of protoplasts exhibiting nuclear localised eYFP varied. 59.2% of mock and 0.005 mM SA-treated protoplasts expressed eYFP, whereas 76.7% of 0.01 mM SA-treated protoplasts expressed eYFP (Fig. 4). Notably, even though the number of eYFP expressing protoplasts was identical in mock and 0.005 mM SA-treated protoplasts, the fluorescence intensity of the individual protoplasts was considerably greater in 0.005 mM SA-treated protoplasts, compared to mock-treated protoplasts (Fig. 4, Fig. 5A). The heterogeneity in nuclear-localised eYFP expression is unlikely to have been caused by temporal differences in the response of individual protoplasts to SA, considering that native eYFP is very stable (Liu and Yoder 2016). Liu and Yoder showed that the half-life of YFP is around 2.5 days in *Medicago truncatula* roots (Liu and Yoder 2016). Therefore, the eYFP expressed during the 24 h protoplast assay was unlikely to be degraded within the timeframe of the experiment. Together, these data suggest (1) that ∼ 60% of the protoplasts are competent to respond to low SA levels and their response strength is SA concentration-dependent, and (2) at higher concentrations of SA a second subpopulation of protoplasts becomes competent to respond to SA. This second subpopulation might be in a more repressed epigenetic state, demanding overall higher SA levels to activate SA-elicited responses. In a tissue context the presence of cells with varying responsiveness to SA might be advantageous. More responsive cells can launch quick local responses, whereas less responsive cells help limit a hypersensitive response to the site of infection.

Separation of eYFP expressing and non-eYFP expressing sub-populations by fluorescence-activated cell-sorting at different SA concentrations could help further elucidate the molecular determinants of protoplast heterogeneity. RNA-seq or RT-qPCR on cell-type specific markers (Kim et al. 2021) of these sub-populations will help identify the various competencies in different cell identities. A combination of CHIP-seq and BS-seq will then allow the determination of the epigenetic states likely dictating the competence of protoplasts of a certain cell identity to respond to different SA concentrations.

### The protoplast assay as a screen for SA-modulating bacteria

Out of ten isolates tested for their potential to modulate PR1 expression, two caused a mild activation with ∼ 1.2× peak fluorescence compared to mock-treated protoplasts and four isolates caused a strong activation with a peak fluorescence of at least ∼ 1.4× the peak fluorescence of mock-treated protoplasts (Fig. 8B). Within the Sphingomonadaceae and Methylobacteriaceae the responses varied notably, with some strains eliciting strong responses and some none. Many *Sphingomonas* protect plants against Pst infection, and the extent of protection varies between different *Sphingomonas* strains (Innerebner et al. 2011). A potential molecular basis for this protection is the activation of SA-related immunity. RNA-seq of plants inoculated with the protective *Sphingomonas melonis* Fr1 strain exhibited gene expression changes indicative of SA-related immune activation, which were absent in the non-protective *Methylobacterium extorquens* PA1 (Vogel et al. 2016). It might thus be that these observed differences in SA response elicitation among the Sphingomonadaceae were linked to the *in planta* protective ability of the strains. The two tested Pseudomonadaceae both showed a strong activation of SA response. This is likely due to their close phylogenetic relationship to the foliar pathogen Pst, and thus an expected high overlap in MAMP profiles. SA-dependent immune responses are known to be effective against biotrophic and hemibiotrophic pathogens, including Pst (Glazebrook 2005). The two Nocardioidaceae (*Rhodococcus* sp. Leaf225 and *Williamsia* sp. Leaf354) elicited a mild activation of SA response (Fig. 8B). Both strains were previously shown to induce fasciation in peas, likely linked to the production of methylated cytokinins (Jameson et al. 2019). Cytokinin appears to enhance plant immunity in a SA-dependent manner and was shown to positively regulate *PR* gene expression (O’Brien and Benková 2013). It is thus likely that the observed increase in fluorescence, as a proxy for PR1 expression, was induced by elevated cytokinin levels. Suppression of plant immunity appears to be a common trait among root-colonising bacteria. Out of 151 tested bacteria 41% were shown to suppress MAMP-triggered root-growth inhibition, a physiological response to the activation of plant immunity (Ma et al. 2021). The observed physiological response appears to be indeed linked to a true suppression of plant immunity rather than a reduction in immune responses by MAMP sequestration, as similar results were observed upon treatment with the DAMP *At*pep1 (Ma et al. 2021). Interestingly, in the current study none of the tested strains suppressed the signal that was observed in mock-treated protoplasts. Whether this is due to differences in leaf- and root-colonising bacteria, or a simple matter of the limited sample set in the current study remains, thus far, unknown.

None of the tested strains caused a fluorescent response in WT protoplasts (Fig. 8A). However, it is possible that such fluorescent responses will be observed in response to some bacteria. In the case of eYFP as a proxy for PR1 expression there are two possible scenarios: (1) Treatment with SA concentrations ≥ 0.05 mM SA led to an eYFP independent increase in fluorescence (Fig. 3A and C), likely caused by protoplast deterioration, as discussed above. Some bacteria potentially elicit a hypersensitivity response in protoplasts and thus cause protoplast-derived autofluorescence. (2) Certain bacteria might produce fluorescent compounds themselves. For example, some Pseudomonads produce fluorescent siderophores, such as the fluorescent pigment pyoverdine, for iron acquisition (Handfield et al. 2000). To test for bacterial autofluorescence we recommend to test bacteria that elicit fluorescence in WT protoplasts, by performing the assay without protoplasts, but in protoplast supernatant, in case bacterial autofluorescence is triggered by plant compounds. In the case of siderophores a potential solution might be pre-treating the bacteria with high concentrations of iron, as high iron levels significantly reduce siderophore production (McRose et al. 2018).

Out of the ten tested strains six originated from arabidopsis leaves, whereas four were derived from leaves of different plants (Ito and Iizuka 1971; Green and Bousfield 1983; Bai et al. 2015; Remus-Emsermann et al. 2016; Schmid et al. 2018). Interestingly, in the selection of strains tested, within the families of Sphingomonadaceae and Methylobacteriaceae the strains that originated from arabidopsis elicited a PR1 response, whereas the strains derived from other plants did not. As plant microbiota are host specific (Knief et al. 2010), it could be that plants adapt to specifically perceive the bacterial strains relevant to them. This would have to be tested with a larger selection of bacterial isolates derived from, and ideally tested on, different plant host species. In support of such potential plant adaptation other studies have shown that flagellin perception is quantitatively different in various plant species and even genotypes (Robatzek et al. 2007; Vetter et al. 2012). Further, these differences in flagellin binding correlate well with the strength of the downstream defence response and bacterial proliferation (Vetter et al. 2012).

## Conclusion

In summary, the protoplast assay is a powerful tool for investigating target gene expression at a large scale. The fluorescence response of healthy protoplasts was in a linear range and can thus be used for a quantitative analysis of the protoplast response to various bacteria. In addition, the measured fluorescence output to SA treatment is consistent with PR1 expression. Further, mock-treated protoplasts expressed eYFP, likely elicited by endogenous SA in response to the protoplast preparation. This allows bidirectional measurements of changes in PR1 expression by bacteria. Depending on the stable transgenic plant lines employed the protoplast assay can theoretically be expanded to any target gene of choice. Measurements can be performed in 96-well plates, allowing for time- and cost-effective large-scale screening efforts.

